# An algebraic approach to parameter optimization in biomolecular bistable systems

**DOI:** 10.1101/045518

**Authors:** Vahid Mardanlou, Elisa Franco

**Affiliations:** Vahid Mardanlou is with Department of Electrical Engineering, University of California Riverside, 900 University Ave., Riverside, CA 92507, USA.; Elisa Franco is with the Department of Mechanical Engineering, University of California Riverside, 900 University Ave., Riverside, CA 92507, USA.

## Abstract

In a synthetic biological network it may often be desirable to maximize or minimize parameters such as reaction rates, fluxes and total concentrations of reagents, while preserving a given dynamic behavior. We consider the problem of parameter optimization in biomolecular bistable circuits. We show that, under some assumptions often satisfied by bistable biological networks, it is possible to derive algebraic conditions on the parameters that determine when bistability occurs. These (global) algebraic conditions can be included as nonlinear constraints in a parameter optimization problem. We derive bistability conditions using Sturm's theorem for Gardner and Collins toggle switch. Then we optimize its nominal parameters to improve switching speed and robustness to a subset of uncertain parameters.

## I. INTRODUCTION

One of the goals of synthetic biology is the design and construction of *de novo* biomolecular systems with behaviors that satisfy user specifications. An important class of functional behaviors is bistability, which is essential to build signaling cascades, developmental networks, and memory elements [1]. Once the general structure of the desired bistable system has been identified, it may be desirable to optimize concentrations and reaction rates of its components to satisfy given performance specifications, while preserving bistability. Because bistability is a complex, global dynamic behavior, the most straightforward option to optimize parameters is by trial and error. To our knowledge, no systematic approaches to this problem are available. (We remark that this challenge is distinct from the problem of parameter optimization in the context of data fitting [2], or network reconstruction [3].)

In this paper we address the problem of defining a systematic procedure to optimize parameters in biomolecular bistable systems: we want this procedure to identify parameters that maximize or minimize a given objective function chosen by the user, while preserving the bistable behavior of the network. For instance, it may be desirable to maximize or minimize the total concentration of some reagents, or to speed up or slow down certain reaction rates based on the specific components used. We formulate our optimization problem so that desired performance is expressed in the objective function, while bistability is guaranteed via the introduction of nonlinear constraints. Nonlinear constraints are obtained from algebraic conditions on the equilibria and on their stability.

Algebraic approaches for optimization have been previously proposed in the context of biochemical networks. In [4], the metabolic flux of a biochemical network is optimized defining an objective function where a single flux, multiplication of fluxes or concentrations of reactants are either maximized or minimized. The network is described using an S-system differential equation model. Algebraic equilibrium conditions are derived and used as constraints within the optimization problem, so that fluxes are optimized while the network is guaranteed to stay in the neighborhood of the desired steady state. In [5], the stationary behavior of a metabolic network is optimized by imposing stability of the equilibrium. The decision variables in this problem can be fluxes, reaction rates or the concentration of some of the reactants. Algebraic equilibrium and stability conditions (derived from a Routh-Hurwitz table) are included in the problem as part of the optimization constraints. The solution of the problem is a stable equilibrium which is optimal with respect to a chosen objective function.

The main limitation of existing approaches is that they consider optimization of *local* (near equilibrium) behaviors. Here, we show that optimization of a network ensuring a *global* behavior like bistability can still be done using an algebraic approach when the system satisfies some structural conditions. Specifically we focus on a class of bistable systems for which the bistability property can be related solely to the number of equilibria of the system and to the sign of the coefficients of the characteristic polynomial. Equilibrium polynomials can be analyzed using Sturm's theorem [6] or a Routh table [7], obtaining a set of inequalities that guarantee a desired number of positive roots. Bistability conditions (global) can thus be expressed as nonlinear constraints to be enforced in the optimization procedure.

The paper is organized as follows. Several notions from algebraic geometry and degree theory are reviewed in Section II; at the end of this section we formulate our optimization problem. Section III describes our case study, for which we derive algebraic conditions for bistability. Results for parameter optimization using our algebraic approach on the described system are presented in Section IV. We summarize and briefly discuss our results in Section V.

## II. BACKGROUND AND PROBLEM FORMULATION

We review definitions and theorems, and formulate assumptions we need to establish algebraic conditions for bistability in biomolecular networks. These conditions are exploited as constraints in the parameter optimization problem described later.

### A. Definitions and assumptions

We consider nonlinear biological dynamical systems that are successfully modeled with ODEs, which may be derived from chemical reactions by applying the law of mass-action or from phenomenological observations. These systems can be modeled as:

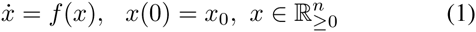

Equilibria (steady states) are states *x* of system (1) satisfying 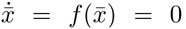. For a system of order *n*, equilibrium conditions are *n* equations in *n* unknowns [8]. If we can merge all equilibrium conditions into a single equation in the form of polynomial function w.r.t. either of the state variables, we refer to it as *master equilibrium condition*.

For every equilibrium point *x̄*, *J*(*x̄*) is the Jacobian of system (1) evaluated at the equilibrium. Polynomial *P*_*x̄*_(λ) = *det*(*λI* – *J*(*x̄*)) = *λ*^*n*^ + *a*_*n*–1_*λ*^*n*–1^ + · · · + *a*_1_λ + *a*_0_ is the characteristic polynomial of the Jacobian evaluated at *x̄*. In many practical cases, the characteristic polynomials of bistable networks have a unique coefficient sign pattern that we summarize in the definition below [6].

#### Definition 1 (F-polynomial)

A polynomial whose coefficients are all non-negative (non-positive) except the constant term ao, which can be either positive or negative, is an F-polynomial. Because the constant term is *a*_0_ = *det*(–*J*(*x̄*)), its sign depends on the determinant of *J*(*x̄*).

This paper focuses on nonlinear systems (1) that are *bistable*.

#### Definition 2 (Bistability)

System (1) is bistable if it presents two asymptotically stable and one unstable equilibrium points.

We consider biomolecular networks whose linearization is a positive system. This property will be used to derive algebraic conditions for bistability.

#### Theorem 1

[9] A linear system *ẋ* = *Ax* is positive if and only if matrix *A* is a Metzler matrix, i.e., its elements satisfy: *a*_*ij*_ ≥ 0, ∀(*i*, *j*) such that *i* ≠ *j*.

Finally, we consider systems that are dissipative. In general, *dissipativity* (see, e.g., [10]) is defined based on the existence of compact, forward invariant subsets of 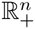 that absorb the system trajectories. The following definition ([11]) is equivalent and easier to verify:

#### Definition 3 (Dissipative system)

A system *ẋ* = *f* (*x*), *x* ∈ ℝ^*n*^ is dissipative if its solutions are eventually uniformly bounded, i.e., there exists a constant *k* > 0 such that:

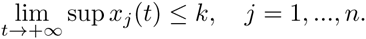

We conclude this section with a list of assumptions we make in the rest of this manuscript.

#### Assumption 1

We assume the linearization of system (1) is a positive system at each equilibrium point.

#### Assumption 2

We assume the characteristic polynomial of system (1) is a F-polynomial at each equilibrium point.

#### Assumption 3

We assume that system (1) is dissipative.

### B. Bistability conditions

The number of equilibria of system (1) can be in many cases established by finding the roots of a master (polynomial) equilibrium condition [6]. The number of positive roots of a polynomial can be established using methods such as Routh table [7] or Sturm’s theorem [6] on the master equilibrium condition of the system. These methods can provide parametric conditions for the presence of a desired number of positive roots, *i.e.* of equilibria. In this paper we foucs on Sturm’s theorem.

*1) Sturm’s theorem approach:* The Sturm sequence associated to a polynomial *p*(*x*) is a set of polynomials ***P*** = {*p*_0_, *p*_1_, …, *p*_*m*_} defined as:

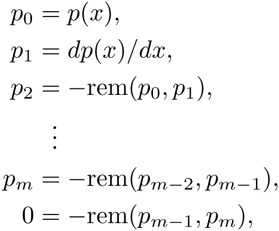

where the symbol rem(*p*_*i*–2_, *p*_*i*–1_) indicates the remainder of the polynomial long division of *p*_*i*–2_ by *p*_*i*–1_ for *i* = 0, 1. It is assumed that *p*_0_ and *p*_1_ are univariate polynomials. The sequence ends when *p*_*m*–1_ divided by *p*_*m*_ gives a remainder of zero.

#### Theorem 2 (Sturm’s theorem)

Let *p*(*x*) be a real-valued univariate polynomial and *a*, *b* ∈ ℝ ∪ {–∞, +∞}, with *a* < *b* and *p*(*a*), *p*(*b*) ≠ 0. Then the number of zeros of *p*(*x*) in the interval (*a*, *b*) is the difference *var*(***P***, *a*) – *var*(***P***, *b*) where *P* is the Sturm sequence of *p*(*x̄*) and the variations *var*(***P***, ***a***) and *var*(***P***, *b*) are the number of times that consecutive nonzero elements of the Sturm sequence — evaluated at *a* and *b*, respectively—have opposite signs.

Sturm’s theorem can thus be used to find conditions for the master equilibrium polynomial to present a desired number of non-negative, real roots by considering the interval [0, ∞).

*2) Algebraic equilibrium conditions are also bistability conditions in a class of bistable networks.:* A bistable system is characterized by the presence of three equilibria; of course, the correct number of equilibria does not in general guarantee bistability.

However, biomolecular bistable networks often satisfy Assumptions 1 – 3; several examples can be found in [6]. For these systems, bistability is guaranteed when three equilibria are present. For the reader’s convenience, we provide additional background on this topic.

#### Theorem 3

Consider a linear positive system *ẋ* = *Ax*, whose characteristic polynomial is an F-polynomial. This system has an unstable equilibrium if and only if the constant term of the characteristic polynomial is negative [7].

#### Definition 4 (Index of an equilibrium point)

The index of a regular equilibrium point *x̄* is the sign of the determinant of *–J* (*x̄*):

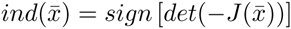

#### Definition 5 (Degree of a system)

The degree of a dynamical system *ẋ* = *f*(*x*), over a set *U* ∈ ℝ^*n*^, having equilibria *x̄*_*i*_, *i* = 1,…, *m*, is defined as:

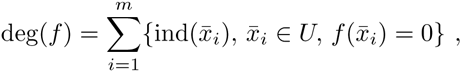

where *x̄*_*i*_ are regular equilibria.

The following theorem is the last result we need to quote:

#### Theorem 4

[11] A dissipative dynamical system *ẋ* = *f*(*x̄*) defined on ℝ^*n*^ has degree +1 with respect to any bounded open set containing all its equilibria [6].

#### Theorem 5

Consider a dynamical system (1), satisfying Assumptions 1–3. If the system has three equilibria, then it is bistable.

#### Proof

Due to Theorem 3, since the index is equal to the sign of the constant term in the characteristic polynomial, a positive index is associated with a stable equilibrium and a negative index is associated with an unstable equilibrium. Due to Theorem 4, we can conclude that when three regular equilibria are present, our systems are bistable.

We conclude remarking that as a consequence of Theorem 5, any algebraic equilibrium conditions that enforce the presence of three equilibria become global algebraic bistability conditions.

### C. Optimization problem

We illustrate the general formulation of our optimization problem. We consider dynamical systems *ẋ* = *f*(*x*,*p*) where p are decision variables (for example, reaction rates), and *p* ∈ Ω. Ω is the state space of all admissible decision variables of the optimization problem.

Our concern is to optimize an objective function ***F***(*p*) which could be a function of a single or a combination of decision variables.

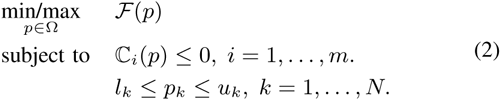

The constraints in the problem above include bistability algebraic conditions, upper and lower bounds for decision variables (*l*_*k*_, *u*_*k*_). Because of Assumptions 1–3, our constraints are independent from the value of the equilibria.

## III. CASESTUDY: GARDNER AND COLLINS TOGGLESWITCH

In this section we introduce the toggle switch by Gardner and Collins [12] and describe the derivation of global algebraic conditions for bistability.

The circuit is composed by two mutually repressing genes, and its nondimensionalized model is:

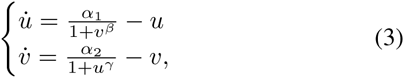

where *α*_*i*_ are the maximal synthesis rates of the i-th repressor, *β* and *γ* are Hill coefficients. The equilibrium conditions of the toggle switch are 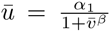 and 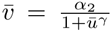 where *ū* and *v̄* are equilibrium points. To simplify our notation, we will denote *ū* and *v̄* as *u* and *v*, dropping the bar. Again for simplicity, we take the value of *β* = *γ* = 2, *i.e.* the repressors are dimers.

*1) Application of Sturm’s theorem to find algebraic equilibrium conditions:* The master equilibrium condition, expressed with respect to variable *υ*, is:

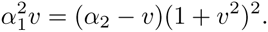

We rewrite it as a fifth order polynomial w.r.t. *υ*,

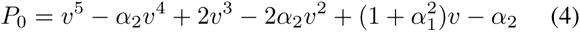

We use Sturm’s theorem to find algebraic conditions guaranteeing that the circuit has exactly three positive roots. First, we find the Sturm sequence *S* = {*P*_0_, · · ·, *P*_5_}, where *P*_*i*_ = *dP*_0_/*dv* and *P*_*i*_ = –*rem*(*P*_*i*–2_, *P*_*i*–1_) for *i* ∈ {2, · · ·, *n*}. Applying Sturm’s theorem, we find 6 possible Sturm sequences where 3 cases for 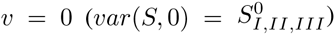 and 2 cases for 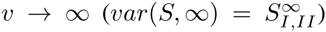 are admissible and yield exactly three positive solutions. These admissible sequences were found with the support of *Wolfram Mathematica*. Furthermore, for *v* → ∞ the sign of *P*_2_ doesn’t have any effect on the total number of sign variations of the sequence 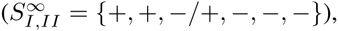, so we are left with 3 distinct cases 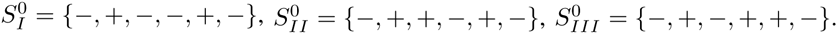.

For the polynomials in the sequence, we find conditions on the parameters so that the appropriate sign changes are obtained. With some tedious derivations (Appendix) we identify a single sufficient condition we can use to guarantee that the master equilibrium polynomial (4) has exactly three positive roots.

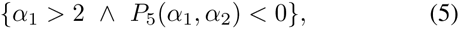

Condition (5) was derived with the support of *Wolfram Mathematica*; details can be found in Appendix A. We will use condition (5) as a constraint of the optimization problem.

In Fig. 1 the mono-stability and bistability regions are plotted using *MATLAB* in a range of *α*_*i*_ ( (0,200] for *i* = 1, 2. In this plot, different colors are associated to the different Sturm sequences 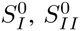 and 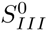.

**Fig. 1.**
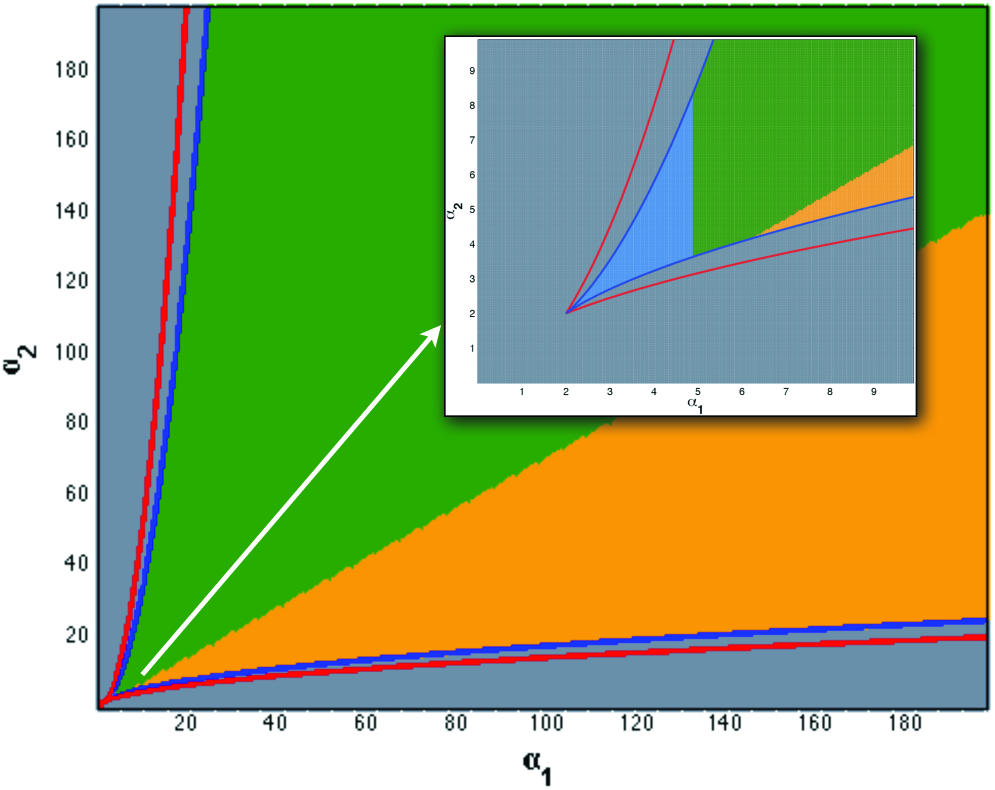
Stability domains of Gardner’s toggle switch. Gray area: monostable domain. Green, light blue, and orange regions are bistability areas. Inset shows details for the bistability region for *α*_1_ ∈ (0; 10]. The green region is associated to Sturm sequence 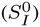, light blue is associated to sequence 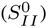 and orange is associated to sequence 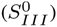. The blue curves are the boundaries of condition (5). The red curves represent the boundaries of conditions *β*_*u*_, *β*_*υ*_ > 2 derived in Section III-3 examining the constant term of the characteristic polynomial.

*2) Verification of Assumptions 1 - 3:*

- Assumption 1: The Jacobian of system (3) is

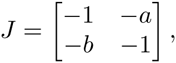

where 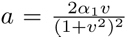 and 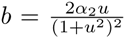 This Jacobian can be transformed into a Metzler matrix by changing sign to its first row and first column. This Assumption is verified regardless of the system’s parameter values.
- Assumption 2: The characteristic polynomial has positive coefficients except the constant term (*a*_0_) which is not sign definite. This Assumption is verified for arbitrary values of parameters.

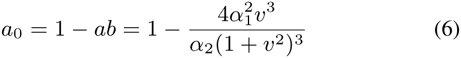 Theorem 3 tells us that the stability of each equilibrium depends on the sign of *a*_0_: the equilibrium is stable if *a*_0_ > 0, and it is unstable if *a*_0_ < 0.
- Assumption 3: It is easy to show that both states of system (3) are eventually uniformly bounded, for arbitrary parameter values. For instance, *u̇* ≤ *a* – *u*, thus *v*(*t*) ≤ max{*v*(0),*αe*^−*t*^}, ∀*t* ≥ 0. *3)Algebraic conditions to determine admissible locations of equilibria:* Theorem 3 indicates that the stability of each equilibrium point is determined by the constant term ao of the characteristic polynomial. We exploit this fact to derive algebraic conditions to determine the location of stable and unstable equilibria. We can rewrite ao as: 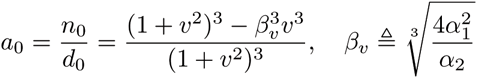 Because the denominator do is positive, the sign of ao is only determined by its numerator no. The sign of no can be studied by rewriting it as:

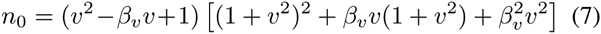 The factor in square brackets is positive, thus the sign of no depends on the sign of *q*(*υ*) = (*υ*^2^ – *β*_*υ*_*υ* + 1), a second order polynomial. Its discriminant 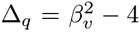 is positive for *β*_*υ*_ > 2 and negative for *β*_*υ*_ < 2. Thus, when *β*_*υ*_ < 2, Δ_*q*_ < 0, *q*(*υ*) > 0 ∀*υ* and the sign of *a*_0_ is always positive in all the domain; we conclude that in this case the system is never bistable, because any existing equilibrium is stable. If *β*_*υ*_ > 2 and Δ_*q*_ > 0, then the sign of *q*(*υ*) changes as follows:

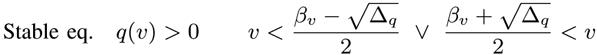

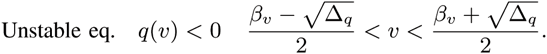 We can derive similar inequalities for variable *u*, by defining the parameter 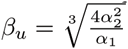. These results indicate that stable equilibria always “sandwich” the only unstable equilibrium, as depicted in Fig. 2. The admissible location of equilibrium points may be a feature we want to automatically optimize. A large distance between the stable equilibria, for instance, means that there is a more significant distinction between the two admissible states; it also means that switching due to perturbations may be more difficult. In contrast, a small distance between stable equilibria may favor rapid toggling between the stable states. We can influence the size of the admissible regions for equilibria by maximizing or minimizing *β*_*υ*_ > 2 and *β*_*u*_ > 2. We note that values of *υ* and *u* for which *n*_0_ = 0 (dashed lines in Fig. 2), and thus *a*_0_ = 0, are not roots of the master equilibrium condition (4) (*Wolfram Mathematica* worksheets are available at [13]). Thus, all admissible equilibria are regular equilibria.

**Fig. 2.**
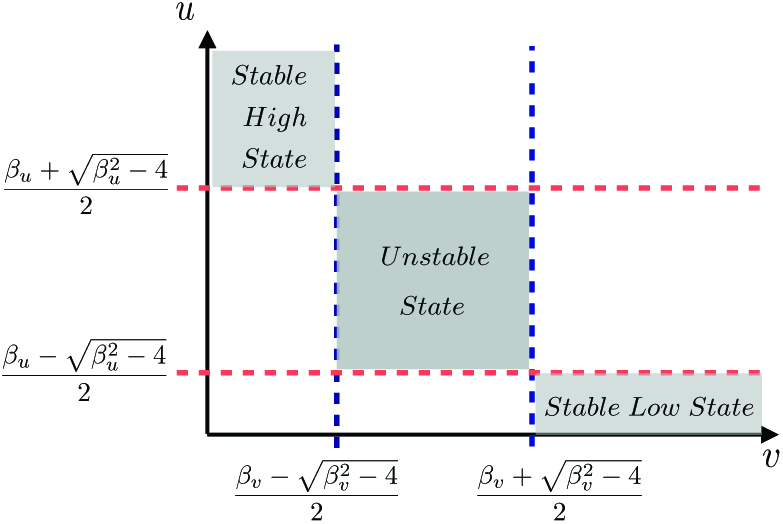
Admissible regions for stable and unstable equilibria determined by analyzing the constant term of the characteristic polynomial.

## IV. PARAMETER OPTIMIZATION EXAMPLES

In this section we formulate the optimization problem on our case study. In this problem, global algebraic conditions for bistability is enforced as part of the optimization constraints as explained in Section II-C. Script was written in MATLAB; optimizations were solved using the genetic algorithm (GA) toolbox. Simulations were done using a 2.13 GHz Intel Core 2 Duo CPU on Linux interface. The length of each single optimization run is 300 seconds on average.

Our goal is to optimize the toggle switch so the admissible regions for its stable steady states are at a minimal distance. If the purpose of the switch is to toggle rapidly, neighboring stable steady states are likely to favor fast switching. As discussed in Section III-3, this can be done by minimizing parameters 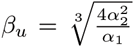 and 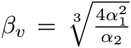In addition, we want to identify nominal values an that are sufficiently far from the boundary of the bistability regions so the system is less sensitive to the perturbations of the parameters: thus we define a “distance” parameter *d*_*i*_ which we want to maximize so that deviations of *α*_*i*_ from the nominal values will still fall in the bistable regions, i.e. 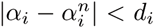. By maximizing di we improve the robustness of the bistable behavior. This goal can be described as:

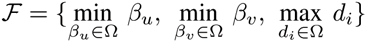

This problem can be converted into a single-objective optimization problem:

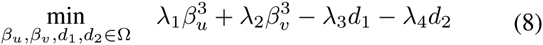

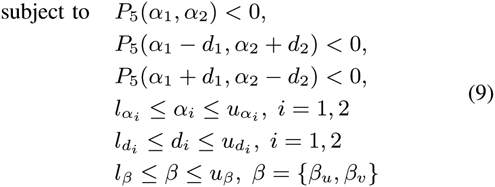

Coefficients *λ*_*i*_*s* in equation (8) are weights used to scale the importance of each parameter in the optimization problem.

For instance, in Fig. 3 we compare changes for the pair *λ*_3_ and *λ*_4_. The i-th rectangle on the right side shows the bistability region w.r.t. the nominal rates 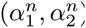 in the center (*c*_*i*_) where the 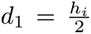 and 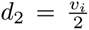. On left side plot in Fig. 3 we have three different cases. In the 1^*st*^ case, the region is a square which means the value for *λ*_3_ equals to *λ*_4_. In this case we consider the robustness issue in the same level for both *α*_*i*_ rates. We can conclude similarly for the other cases. Here, *β*_*u*_, *β*_*υ*_ are combinations of parameters, thus can be considered auxiliary variables (see Section II-C).

**Fig. 3.**
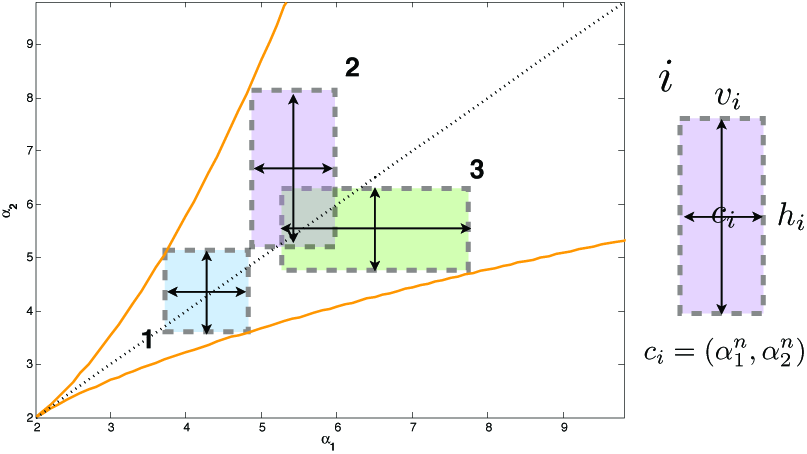
Effects of the weighting coefficients *λ*_3_ and *λ*_4_ in objective function (8).

Variables are scaled to be comparable; while both *α*_*i*_s have same dimensions and operational ranges as well as *d*_*i*_s, *β*_*u*_ and *β*_*υ*_ are scaled to their third power to be comparable with the other variables.

Regarding the constraints (9), they include the bistability condition (5) which must be valid for the new coordinates (*α*_1_ ∓, *d*_1_, *α*_2_±*d*_2_). These coordinates are the diagonal vertices of the rectangles surrounding the nominal rates in Fig. 3. The last three inequalities in (9) are the boundary constraints on the parameters of the problem.

Fig. 4 shows solutions of the optimization problem found using *MATLAB GA Optimization Toolbox*. We ran 350 simulations with randomly selected initial points (uniformly sampled). Each dot corresponds to a local minimum of a single run of the GA after 150 generations; colors from blue to red indicate an increasing value of the objective function. Dots with the same color on the different panels of Fig. 4 correspond to the same optimization round. We note that the optimization results are clustered in parameter space; choosing nominal parameters at the center of these clusters would guarantee a robust performance in the presence of uncertainty.

**Fig. 4.**
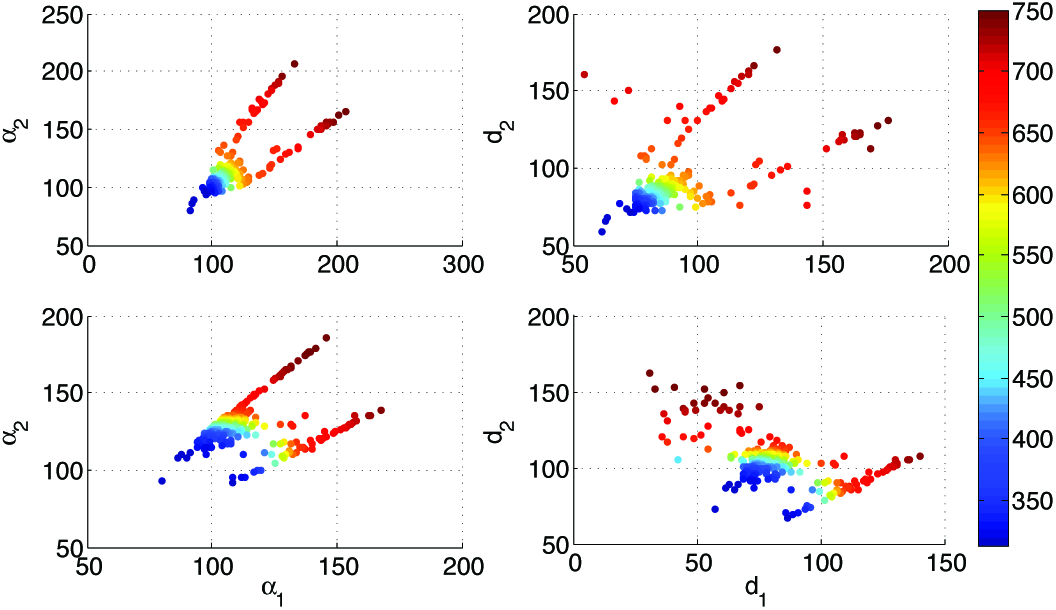
The Genetic Algorithm optimization result for 350 uniformly random selected initial points after 150 generations. (Left): *α*_2_ – *α*_1_ plots. (Right): *d*_2_ – *d*_1_ plots. (Upper): ∧ = (*λ*_1_,*λ*_2_,…) = (0.5, 0.5,0.1,0.1) and (Bottom): ∧ = (0.5,0.5, 0.1, 0.4).

## v. CONCLUSIONS

We have described an approach to parameter optimization for bistable biological networks. We showed that global bistability conditions derived from standard algebraic methods can be included as nonlinear constraints in the optimization procedure. These conditions can be derived for systems that satisfy a series of assumptions: dissipativity and positivity are relatively mild assumptions in biomolecular bistable networks, and are often structural properties [7] that do not depend on the parameters. Assumption 2, which prescribes the positivity of all coefficients of the characteristic polynomial, may limit the applicability of our approach, but several biomolecular networks in the literature are found to satisfy this requirement [6]. Clearly, algebraic tools such as Sturm’s theorem can become cumbersome to apply to systems of large order. However, this method is applicable to bistable modules within larger networks, in which inputs or interconnections with the large networks may be regarded as parameters of the circuit.

## APPENDIX

From the Sturm sequence polynomials, we isolated terms that determine the sign of the sequence element (as a function of *α*_1_ and *α*_2_). Term 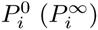, for instance, is the term that determines the sign of *P*_*i*_(*υ* → 0) (*P*_*i*_(*υ* → ∞)). Due to space limitations, we do not report the full expressions for the Sturm sequence polynomials.

In the following sections the symbol “∨” means “*or*” and symbol “∧” means “*and*”.

- *P*_0_(*υ* → 0) = –*α*_2_ < 0, ∀*α*_2_ > 0.
- *P*_0_(*υ* → ∞) → ∞ > 0.
- 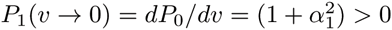
- *P*_1_ (*υ* → ∞) → ∞ > 0.

*P*_0_ and *P*_1_ are sign definite regardless of the values of *α*_1_ and *α*_2_.

- *P*_2_(*υ* → 0): its sign is determined by 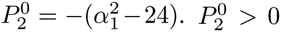 when 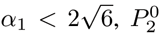 when 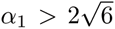. The sign of *P*_2_(*υ* → 0) does not depend on *α*_2_.
- *P*_2_(*υ* → ∞): Its sign is determined by 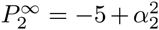. 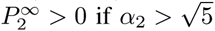 and 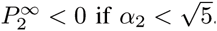
- *P*_3_(*υ* → 0). The sign of *P*_3_ is determined by:

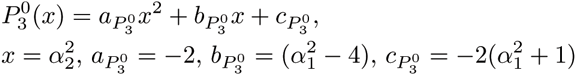 The discriminant is 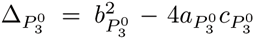 and we define 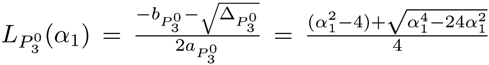 and 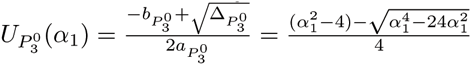.

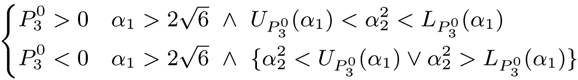 For 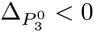 we find that 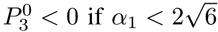.
- *P*_3_(*υ* → ∞): its sign is determined by 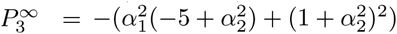

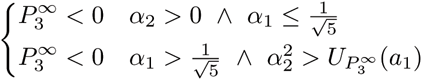

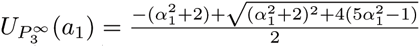
- *P*_4_(*υ* → 0). The sign of *P*_4_ is determined by:

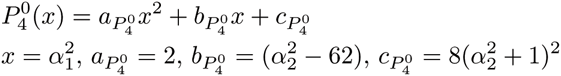 If 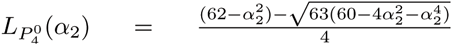 and 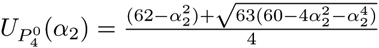 then, 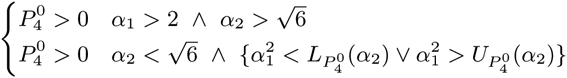
- *P*_4_(*υ* → ∞). The sign of this term is determined by

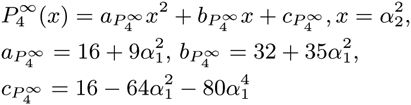 Then we define 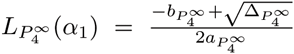 where 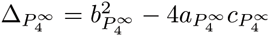 The conditions on the sign of 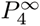 are:

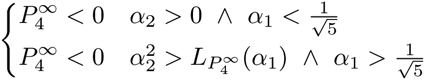
- *P*_5_(*u* → 0, ∞): their sign is determined by:

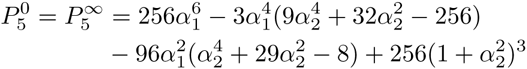 The last polynomial is *P*_5_(*υ* → ∞) = *P5*(*υ* → 0) < if *α*_1_ > 2 and 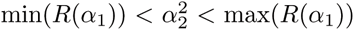. The *R*(*α*_1_) is the set of positive real roots of the polynomial 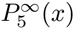 where 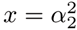.

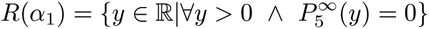 It is easy to prove that the aforementioned polynomial has a single negative real root or 2 positive and 1 negative real roots depending on the value of *α*_1_.

